# Stem cell-laden hydrogel bioink for generation of high resolution and fidelity engineered tissues with complex geometries

**DOI:** 10.1101/2021.04.15.439978

**Authors:** Oju Jeon, Yu Bin Lee, Sang Jin Lee, Nazilya Guliyeva, Joanna Lee, Eben Alsberg

## Abstract

Recently, 3D bioprinting has been explored as a promising technology for biomedical applications with the potential to create complex structures with precise features. Cell encapsulated hydrogels composed of materials such as gelatin, collagen, hyaluronic acid, alginate and polyethylene glycol have been widely used as bioinks for 3D bioprinting. However, since most hydrogel-based bioinks may not allow rapid stabilization immediately after 3D bioprinting, achieving high resolution and fidelity to the intended architecture is a common challenge in 3D bioprinting of hydrogels. In this study, we have utilized shear-thinning and self-healing ionically crosslinked oxidized and methacrylated alginates (OMAs) as a bioink, which can be rapidly gelled by its self-healing property after bioprinting and further stabilized via secondary crosslinking. It was successfully demonstrated that stem cell-laden calcium-crosslinked OMA hydrogels can be bioprinted into complicated 3D tissue structures with both high resolution and fidelity. Additional photocrosslinking enables long-term culture of 3D bioprinted constructs for formation of functional tissue by differentiation of encapsulated human mesenchymal stem cells.

## Introduction

Hydrogels are three-dimensional (3D) networks of hydrophilic polymers that can hold a large amount of water in their swollen state,^1, 2^ and over the past few decades, they have been widely used in the tissue engineering and regenerative medicine fields due to their excellent biocompatibility^3, 4^ and capacity to be engineered to mimic aspects of the native extracellular matrix (ECM) of tissues. A variety of hydrogels, endowed with special characteristics, such as tunable physical and biochemical properties, stimuli responsiveness, multiple-network crosslinking and ultra-toughness, have been adopted with innovative strategies, like photolithography, interpenetrating polymer network formation, coacervation and microgel assembly, for biomedical engineering applications.^5, 6, 7, 8, 9^ However, they are limited in their ability to be used to create precisely controlled complex structures and intricate 3D microarchitectures replicating those in tissues and organs.^10^ Recently, 3D bioprinting has been explored as a promising technology for creating engineered tissue constructs with complex structures at high resolution.^11^ Cell encapsulated hydrogels composed of materials such as gelatin, collagen, hyaluronic acid, alginate and polyethylene glycol have been used extensively as bioinks for 3D printing.^11, 12, 13^ Since most hydrogel-based bioinks do not allow rapid stabilization immediately after extrusion printing, achieving high resolution and fidelity and self-supporting tissue constructs remains a significant challenge in 3D hydrogel bioprinting.^14, 15^ Thus, there is a critical need to develop new hydrogel-based bioinks that can be applied to directly print structures of clinically relevant size-scale with high printing resolution and fidelity, tunable mechanical strength and degradation rates, and high cell viability.^16^

To overcome this limited capacity for rapid mechanical stabilization of hydrogel-based bioinks, multi-material, interpenetrating network, nanocomposite, supramolecular and shear-thinning material-based bioinks ^16, 17^ and supporting medium systems composed of microgel slurries, cell aggregates and shear-thinning hydrogels^14, 18, 19, 20, 21^ have been developed to improve hydrogel printability and mechanical support for 3D printed structures. Among these, shear-thinning hydrogels are often favored since they undergo a substantial reduction in viscosity under increasing shear stress.^22^ Some shear-thinning hydrogels exhibit viscous flow under shear and self-healing upon removal of applied shear stress.^23^ However, many of these shear-thinning bioinks rely on nonspecific interactions between macromers or the development of bioinks by modifications enabling long-range interactions between specific binding molecules on macromers, resulting in prolonged self-recovery times following shear-thinning after extrusion through a needle.^24, 25^ This may limit their suitability as bioinks because they may collapse due to instability before self-healing, which results in the poor resolution and fidelity of 3D printed structures.^26^ To achieve rapid self-healing of hydrogels, supramolecular hydrogels have been developed as shear-thinning and self-healing hydrogel bioinks, which show rapid self-healing time, based on guest-host interactions.^27^ Though due to steric hindrance effects of long macromer chains, the guest-host complexation can be retarded because the host molecules cannot efficiently interact with the guest molecules.^28, 29^ Moreover, non-covalent crosslinking of the guest-host molecules endows the hydrogels with poor mechanical properties.^30^

Here, ionically crosslinked oxidized and methacrylated alginate (OMA) has been utilized as a bioink, which can be further stabilized after bioprinting via secondary crosslinking (**Figure 1** and Figure S1). The OMA bioink exhibits shear-thinning and rapid self-healing properties (**Figure 2**, Figure S2 and Figure S7), and are expected to be applicable to 3D bioprinting systems with high resolution and fidelity. While OMA bioinks could be directly extruded through the printing needle as continuous fibers via their shear-thinning properties, the 3D printed OMA constructs could be stabilized and maintained through rapid self-healing properties. We have successfully demonstrated that stem cell-laden calcium-crosslinked OMA hydrogels can be bioprinted into complicated 3D tissue structures with high resolution and fidelity. Additional photocrosslinking enables long-term culture of 3D bioprinted constructs for formation of functional tissue by differentiation of encapsulated human mesenchymal stem cells (hMSCs).

**Figure 1.**
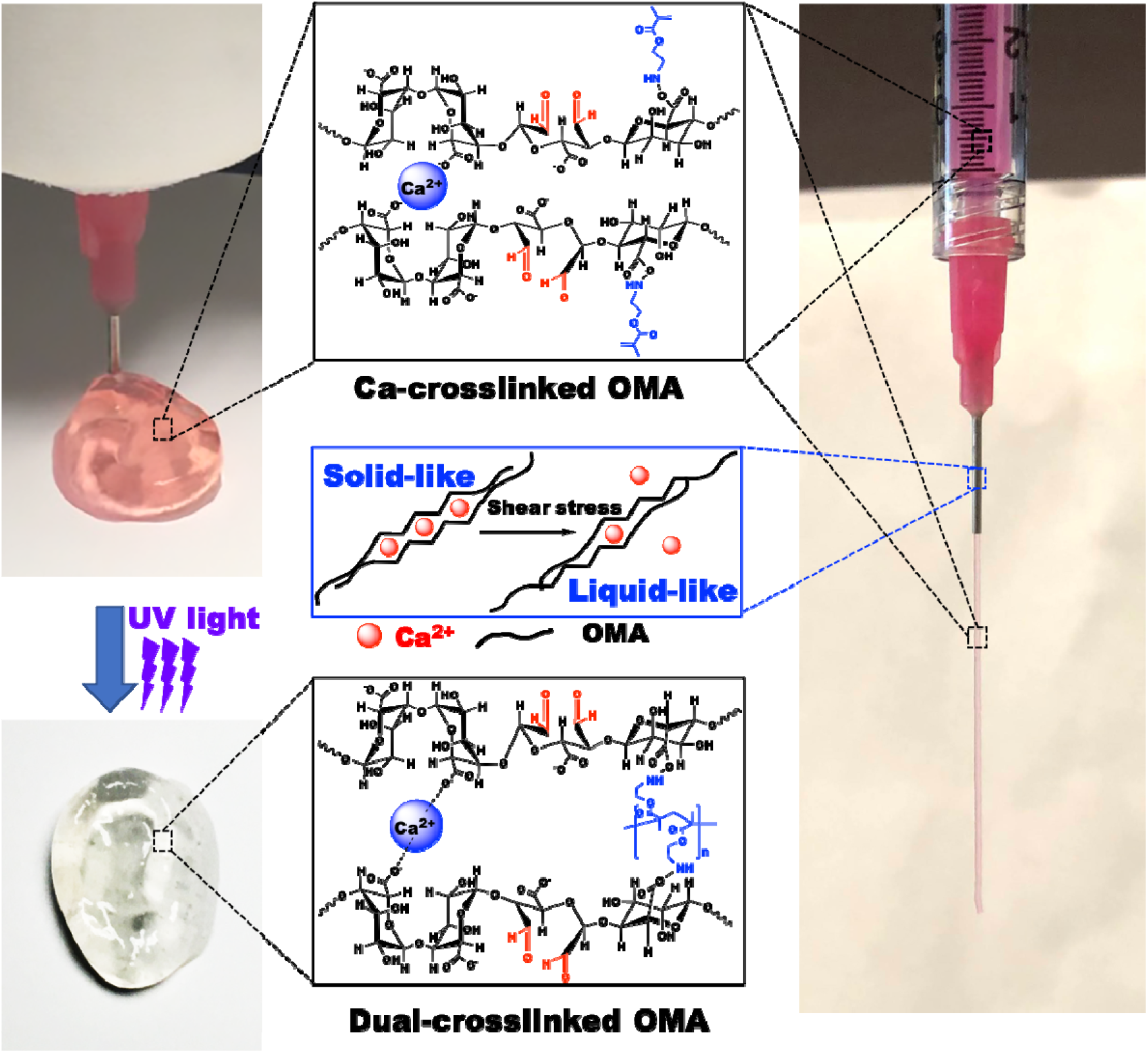
Schematic illustration of photocrosslinkable calcium-crosslinked OMA bioink. The 3D bioprinted OMA bioink could be further photocured to produce a chemically and mechanically stable biomimetic 3D bioprinted construct.

## Results and Discussion

3D bioprinting is an important tool for the development of complex structures of tissues and organs for tissue engineering and regenerative medicine applications.^2^ Since the hydrogel-based bioinks for 3D bioprinting are extruded via syringe through a narrow printing nozzle, they must possess a balance between a high viscosity for rapid gelation after extrusion and low shear stress for cytocompatibility, which is difficult to achieve at the same time in a biomaterial.^31^ Since the shear-thinning and self-healing characteristics and low shear yield stress could enhance the 3D printing capacity of bioinks,^22^ OMA bioinks were evaluated to determine whether they exhibit these properties before utilizing them for 3D bioprinting. The ionically crosslinked OMA bioinks exhibited shear-thinning behavior, characterized by the decrease in viscosity with increasing shear rate [Figure 2(A)], while the uncrosslinked OMA solution (OMA-0) exhibited typical liquid behavior, indicating significantly lower and variable viscosities over the shear rate screen (Figure S2(A)). Additionally, all of the OMA bioinks exhibited low shear yield stress [Figure 2(B) and Figure S2(B), <5 Pa], which was determined using shear stress sweeps, indicating that only a small stress is required to allow the OMA bioinks to flow. Frequency sweep tests of the OMA bioinks showed significantly higher G’ than G”, indicating that the OMA bioinks were mechanically stable [Figure 2(C-F) and Figure S3(A-D)], while the uncrosslinked OMA solutions show typical viscoelastic liquid behavior (Figure S4). In oscillatory strain sweep tests, G” of the OMA bioinks surpassed G’ at approximately 25 % strain [Figure 2(G-J) and Figure S5(A-D)], indicating the phase change from solid-like to liquid-like, which is an important property of hydrogel materials for injectability and/or printability through a printing nozzle. In contrast, the uncrosslinked OMA solutions show typical viscoelastic liquid behavior [Figure S6(A-B)]. When investigated under the cyclic strain sweeps by alternating low (1%) and high (100%) strains, the OMA bioinks went from solid-like to liquid-like behavior in response to strain [Figure 2(K-N) and Figure S7(A-D)]. Furthermore, the responses of shear moduli to high strain and recoveries at low strain were rapid and repeatable. The combination of shear-thinning, low shear-yielding and self-healing properties allows for the rapid transition from solid-like to liquid-like behavior.^32^ Moreover, the rapid recovery of mechanical properties after removal of shear stress, as occurs after the deposition of the OMA bioinks, enables their stabilization immediately after extrusion. These properties make the OMA bioinks well-suited for injection and extrusion-based 3D bioprinting. These characteristics were further confirmed with various OMA bioinks prepare with low viscosity alginate (Figure S8-S12). OMA-20, OMA-25 and OMA-40 bioinks with different amounts of calcium exhibited similar high fidelity to the printed structure of a cuboid, while the OMA-15 bioink exhibited the lowest fidelity of the 3D printed structure [Figure 2(O) and Figure S13]. Therefore, the OMA-20 bioink (high viscosity 1OX20MA; 1 and 20 % theoretical oxidation and methacrylation, respectively), which had the lowest amount of calcium ions and the crossover point of G’ and G” at the lowest shear strain [Figure 2(H)] amongst the compositions which printed at high fidelity, was used for subsequent experiments.

**Figure 2.**
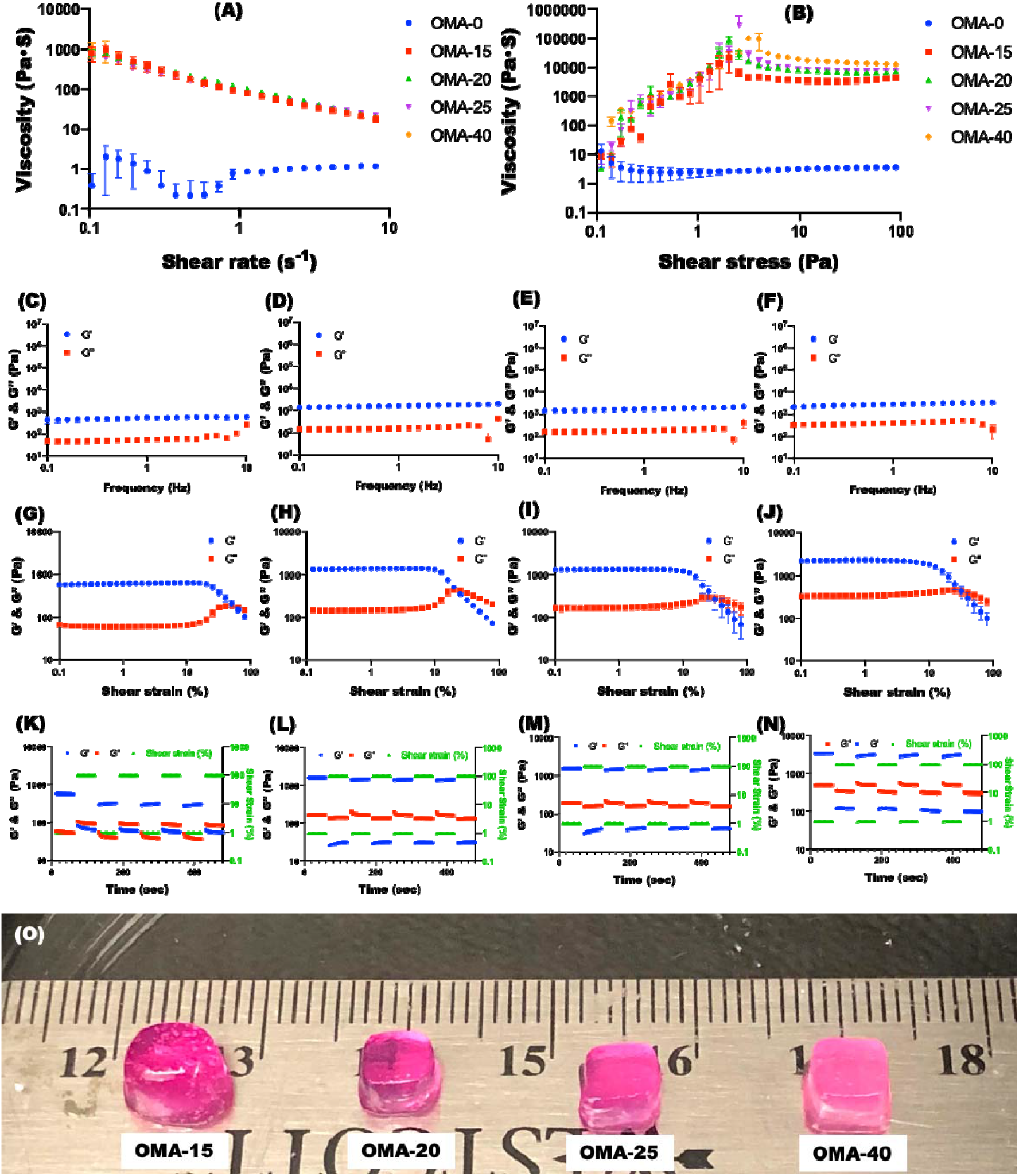
Characterization of OMA (1OX20MA) bioinks synthesized from high viscosity alginate. Viscosity measurements of the OMA bioink as a function of **(A)** shear rate and **(B)** shear stress demonstrate its shear-thinning and shear-yielding behaviors, respectively. Frequency sweep tests of **(C)** OMA-15 [OMA+ 15 μl CaSO4 (1.22 M)], **(D)** OMA-20 [OMA+ 20 μl CaSO4 (1.22 M)], **(E)** OMA-25 [OMA+ 25 μl CaSO4 (1.22 M)], and **(F)** OMA-40 [OMA+ 40 μl CaSO4 (1.22 M)] bioinks indicate that the OMA bioinks were mechanically stable. Strain sweep tests of **(G)** OMA-15, **(H)** OMA-20, **(I)** OMA-25, and **(J)** OMA-40 bioinks. *G* and *G”* crossover of the OMA bioinks as a function of shear strain exhibit their gel-to-sol transition at higher shear strain. Shear moduli changes during dynamic strain tests of **(K)** OMA-15, **(L)** OMA-20, **(M)** OMA-25, and **(N)** OMA-40 bioinks with alternating low (1%) and high (100%) strains at 1 Hz demonstrate their rapid transitions between solid-like and liquid-like behavior within seconds, which indicates self-healing or thixotropic properties. (O) Photograph of the 3D printed structures (5×5×3 mm) using the OMA bioinks on the BioX printer.

To mimic native tissue architecture using an extrusion-based 3D bioprinting, it is important to evaluate the resolution of bioinks. Since the diameter of a printing nozzle has a significant effect on the resolution of printed constructs,^33^ the effect of the printing needle diameter on bioprinting resolution was investigated. The robustness of high-resolution printing was determined by measuring the diameter of printed OMA filaments with various printing needle gauges from 22 to 27 G [**Figure 3(A)**]. As the inner diameter (ID) of the printing needles decreased from 413 μm (22 G) to 210 μm (27 G), the diameters of printed OMA filaments decreased from 361 μm to 223 μm [Figure 3(B)], which were 88 to 106 % of the inner diameters of the needles, respectively, and confirmed the capability of high-resolution printing. This trend was further confirmed with OMA bioinks prepared with low viscosity alginate. 3D printed OMA filaments with 22 G need exhibited higher solution compare to 3D printed filaments with 20 G needle (Figures S14).

**Figure 3.**
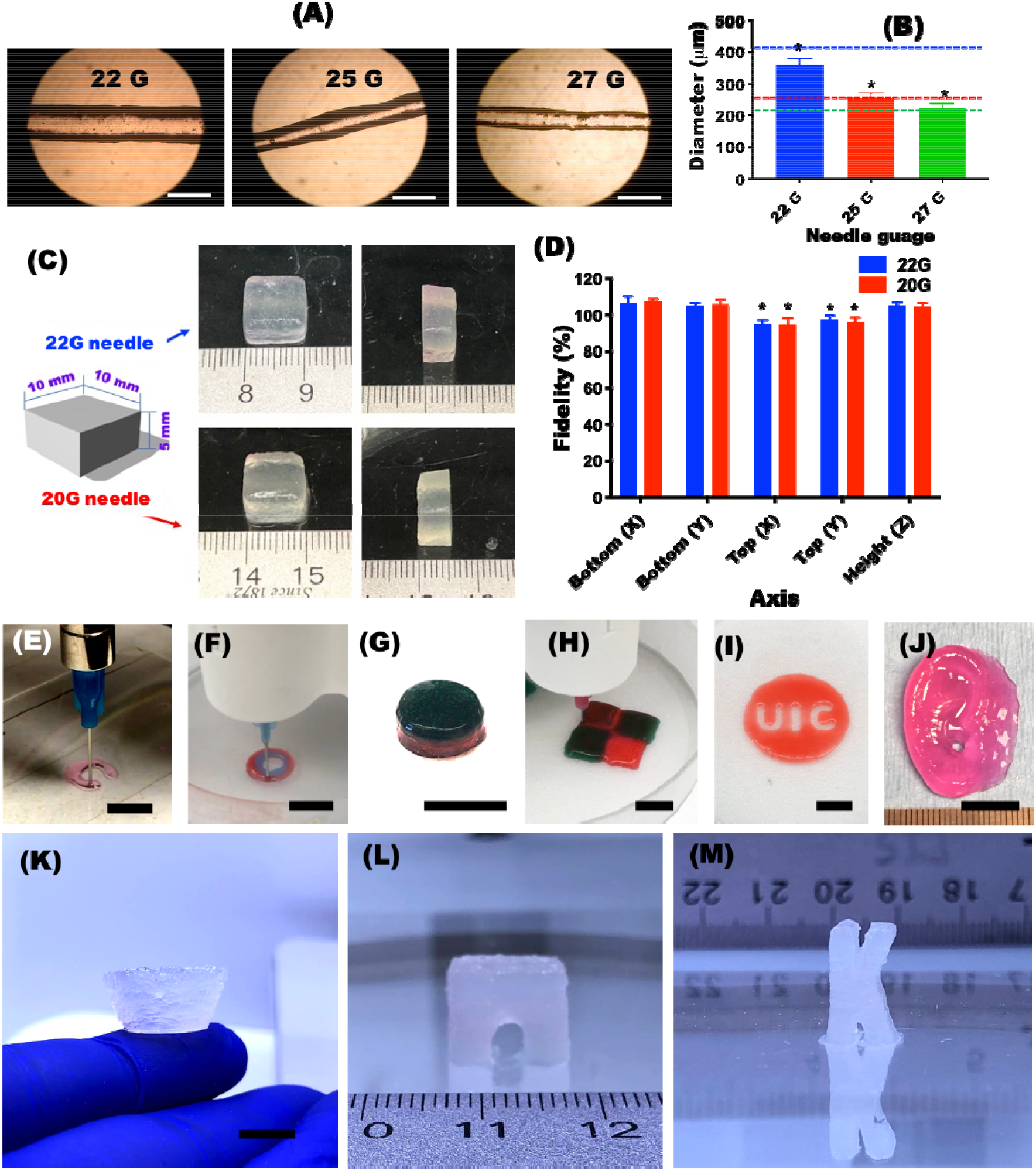
**(A)** Images of 3D filaments printed in a straight line using the modified Printrbot printer with OMA bioinks synthesized from high viscosity alginate and 22 G (ID=410 μm), 25 G (ID=260 μm), and 27 G (ID=210 μm) printing needles and **(B)** their mean diameters. Colored dotted lines indicate the inner diameter of each respective printing needle. Scale bars indicate 500 μm. *p<0.05 compared to other groups. Fidelity of the 3D printed structures with various printing needle sizes. **(C)** Images of 3D printed structures using the modified Printrbot printer with 22 G and 20 G printing needles, **(D)** their quantified fidelity, demonstrating high fidelity of the 3D printed structures. Images of 3D printed structures of **(E)** a letter “C” formed using the modified Printrbot printer, **(F)** a concentric-ring fabricated using the Biobot printer, **(G)** a two-phase cylinder formed using the BioX printer, **(H)** a checkerboard-patterned structure fabricated using the Biobot printer **(I)** the UIC logo formed using the BioX printer and **(J)** an ear formed using the modified Printrbot printer. Images of 3D printed overhang geometries of **(K)** a bowl, **(L)** a bridge and **(M)** a letter “K” using the BioX printer with OMA bioinks synthesized from high viscosity alginate and 22 G printing nozzles. The black scale bars indicate 1 cm.

One of the primary goals of 3D bioprinting is to print complex geometries. However, current hydrogel-based bioinks lack the mechanical strength as well as printability to print macroscale constructs with high fidelity.^32^ Although high resolution of hydrogel bioink filaments have been reported in some studies,^26^ there are still limitations with respect to achieving high shape fidelity of 3D printed structures. To demonstrate the OMA bioinks’ ability to print macroscale shapes with high shape fidelity, cuboids (10×10×5 mm) were printed with 20 (ID=603 μm) and 22 G (ID=413 μm) needles [Figure 3(C) and Figure S15], measured the dimensions of the 3D printed structures (X, Y and Z axes) and then calculated the fidelity (%) by comparison with the 3D digital images. The quantified fidelities of the 3D printed structures were 92 to 110 % [Figure 3(D)], indicating the capability of high fidelity, and there was no significant difference in fidelity achieved with the 20 and 22 G printing needles. Furthermore, the OMA bioinks exhibited high fidelity in forming complex and large-scale structures, such as a letter, a concentric-ring, a two-phase cylinder, a patterned structure, the UIC logo [Figure 3(E-J)], and an ear [Figure S16]. Although complex geometries have been reported in a number of studies using hydrogel bioinks, it is currently still very challenging to 3D extrusion print hydrogel bioinks in overhang geometries without additional supporting structures or materials due to their mechanical instability. Instantaneous recovery of bioinks’ mechanical strength after extrusion is essential to prevent collapse of overhang layers.^34, 35^ Since our OMA bioinks are mechanically stable and exhibit rapid self-healing (Figure 2, S1, S3 and S7), various overhang geometries with self-supporting structural integrity could be extrusion printed using the OMA bioinks without supporting devices or materials [Figure 3(K-M) and Figure S17], which to our knowledge has never been reported before.

Since cell-laden OMA bioinks dispensed from a nozzle are exposed to shear stress and additionally low-level UV light during photocrosslinking to further stabilize the 3D printed constructs, the cytocompability of the bioprinting process was analyzed by measuring cell viability after the printing process and photocrosslinking. High cell viability was observed after printing process and photocrosslinking, indicating that the macromers, the bioprinting process and the photocrosslinking were all cytocompatible [**Figure 4(A-C)**].

**Figure 4.**
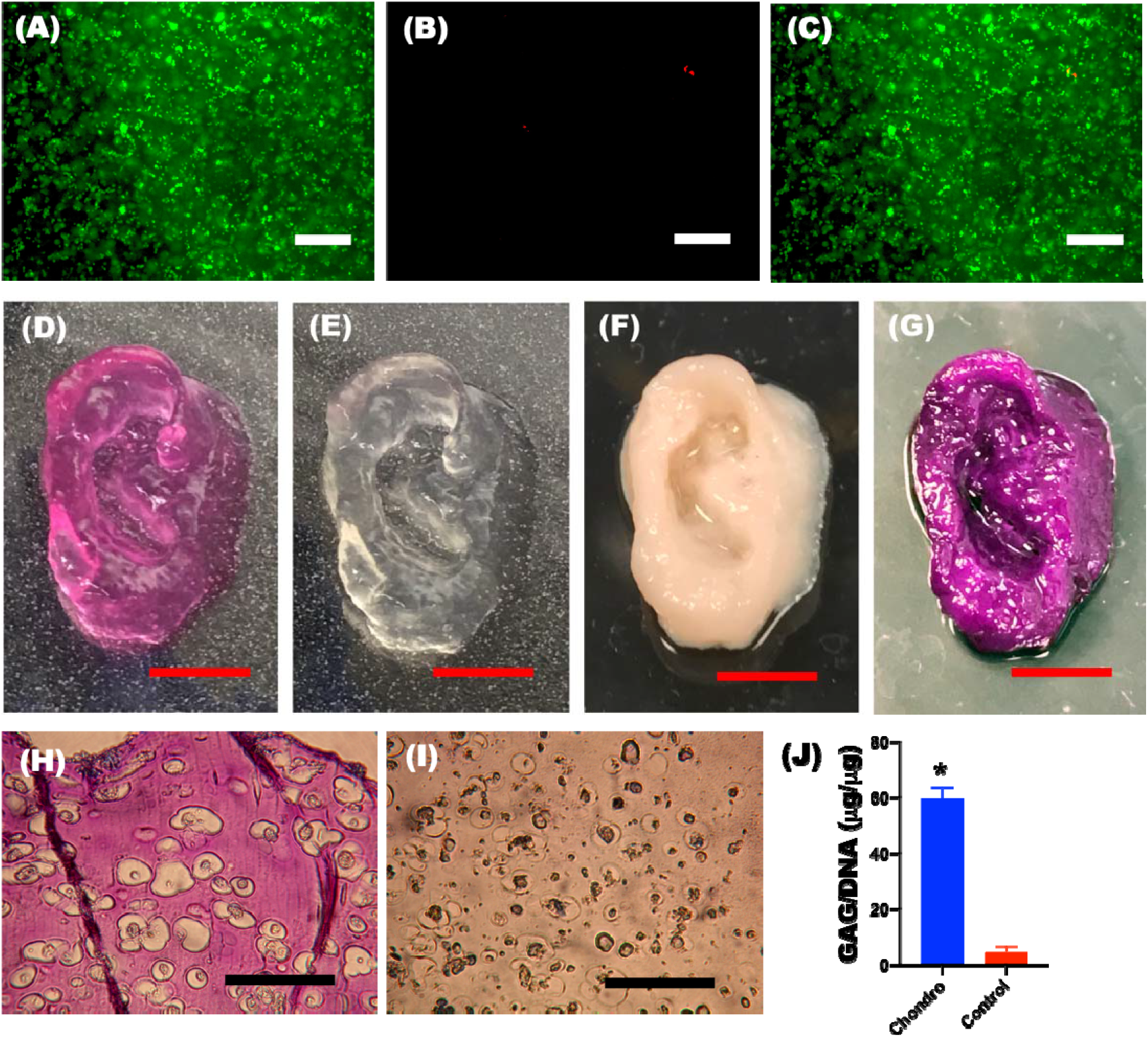
Differentiation of 3D bioprinted constructs using the modified Printrbot printer with hMSC-laden OMA bioinks synthesized from high viscosity alginate. Representative **(A)** live, **(B)** dead and **(C)** merged photomicrographs of photocrosslinked 3D printed constructs at day 0. 3D printed ear using hMSC-laden OMA bioink **(D)** before and **(E)** after photocrosslinking. Chondrogenically differentiated 3D printed ear **(F)** before and **(G)** after Toluidine blue O staining. Photomicrographs of Toluidine blue O stained construct sections cultured in **(H)** chondrogenic and **(I)** growth media. **(J)** Quantification of GAG/DNA in the 3D printed constructs. White, red and black scale bars indicate 200 μm, 1 cm and 100 μm, respectively. *p<0.05 compared to Control.

After validating the printability and cytocompability of the OMA bioinks, the hMSC-laden constructs were printed [Figure 4(D)] for long-term culture to investigate cartilage tissue formation by chondrogenic differentiation of hMSCs. Photocrosslinked 3D bioprinted ears [Figure 4(E)] and cuboids were cultured for 4 weeks in chondrogenic differentiation media, and after 4 weeks, chondrogenically differentiated tissue constructs were harvested [Figure 4(F)]. Differentiation down the chondrogenic lineage and resultant formation of cartilage tissue were confirmed via Toluidine blue O staining; intense purple color was observed throughout the ear constructs [Figure 4(G)] and sectioned cuboid samples [Figure 4(H)], while there was only slight light blue color, which is non-positive staining for GAG, on the 3D printed whole ear (Figure S18) and no positive staining of the sectioned 3D printed cuboid constructs [Figure 4(I)] cultured in growth media. Lacunae structures were also observed in sectioned slides of chondrogenically differentiated constructs [Figure 4(H)], indicating maturation of cartilage tissues. Successful tissue formation by the 3D printed hMSC-laden OMA bioinks was further confirmed by quantification of cartilage tissue glycosaminoglycan (GAG) production in the cuboids [Figure 4(J) and Figure S19].

Numerous natural hydrogel-based cell-laden bioinks have been reported in efforts to biofabricate tissues and organs in the tissue engineering and regenerative medicine fields. However, it’s challenging to achieve high shape fidelity in 3D printed constructs of clinically relevant sizes using hydrogel bioinks due to their aforementioned mechanical instability.^32, 36^ Furthermore, the successful clinical application of cell-laden 3D printed hydrogel constructs depends on the ability to modulate their biochemical and physical properties to create hierarchically complex microenvironments that can regulate encapsulated cell behaviors, such as proliferation, differentiation, migration and apoptosis.^37, 38, 39^ Therefore, development of bioinks that permit tuning of these properties is essential to tackle current challenges of hydrogel-based bioinks and regenerate biomimetic functional tissue constructs.^37^ Not only does the bioink system reported here address the problems of fidelity, resolution and mechanical stability, but it is possible to independently control the OMA biochemical (e.g., cell adhesivity) and physical (e.g., mechanical, swelling and degradation profile) properties and bioactive molecule delivery capacity.^1, 6, 40, 41, 42, 43, 44, 45, 46^ In this respect, OMA bioinks provide critical unique advantages over the other hydrogel-based bioinks, including the ability to guide the regeneration of tissues of clinically relevant sizes by 3D printing constructs using bioinks with tissue specific cues to guide the function of encapsulated cells. In addition, the OMA bioinks allow controlled deposition of patterned, multiple layered structures and complex geometries such as overhangs with high shape fidelity to mimic the sophisticated and heterogeneous architectures of developing and native tissues without supporting materials, such as microgel slurry baths or thermoplastics.^18, 47^ Therefore, platform technology is a promising candidate to achieve the regeneration of complex functional 3D tissues and organs, which require precise microarchitectures comprised of multiple types of cells and ECMs in precise spatial distributions.^48^

In conclusion, photocrosslinkable OMA bioinks have been prepared using an ionically crosslinked OMA hydrogels to create complex 3D tissue structures. The OMA bioinks exhibited both high resolution of 3D printed filaments and high fidelity of 3D printed structures. Importantly, to our knowledge, this is the first time complex overhang geometries could be extrusion printed with bioinks without supporting devices or materials. The 3D printed constructs were further photocrosslinked for increased stability. When hMSCs were incorporated in the OMA bioinks, their viability was high after the 3D printing process and subsequent photocrosslinking. Additionally, 3D printed constructs using hMSC-laden OMA bioinks could be successfully induced to formed cartilage tissue. 3D printing of stem cell-laden OMA bioinks provides a powerful and highly scalable platform technology for 3D tissue construction. Further, the applications of printed constructs using this approach can be easily expanded to include the incorporation of other cell types and bioactive factors to engineer other tissues of interest.

## Methods

### Synthesis of OMA

The dual-crosslinkable oxidized and methacrylated alginate (OMA) was prepared by the oxidation and methacrylation of alginates.^21, 49^ Briefly, sodium alginate (10 g, Protanal LF120M [high viscosity (251 mPa•S at 1 % water) or low viscosity (157 mPa•S at 1% water), FMC Biopolymer] was dissolved in ultrapure deionized water (diH_2_O, 900 ml) overnight. Sodium periodate (0.1, 0.2 or 0.5 g, Sigma) was dissolved in 100 ml diH_2_O, added to the alginate solution under stirring in the dark at room temperature (RT) and then allowed to react for 24 hrs. To synthesize OMA, 2-morpholinoethanesulfonic acid (MES, 19.52 g, Sigma) and NaCl (17.53 g) were directly added to an oxidized alginate (OA) solution (1 L), and the pH was adjusted to 6.5 with 5 N NaOH. N-hydroxysuccinimide (NHS, 2.12 g; Sigma) and 1-ethyl-3-(3-dimethylaminopropyl)-carbodiimide hydrochloride (EDC, 7.00 g; Sigma) (molar ratio of NHS:EDC = 1:2) were added to the mixture to activate 20% of the carboxylic acid groups of the alginate. After 5 min, 2-aminoethyl methacrylate (AEMA, 3.04 g, Polysciences) (molar ratio of NHS:EDC:AEMA = 1:2:1) was added to the product, and the reaction was maintained in the dark at RT for 24 hrs. The reaction mixture was precipitated with the addition of acetone in excess, dried in a fume hood, and rehydrated to a 1 w/v % solution in diH_2_O for further purification. The OMA was purified by dialysis against diH_2_O (MWCO 3500, Spectrum Laboratories Inc.) for 3 days, treated with activated charcoal (5 g/L, 100 mesh, Oakwood Chemical) for 30 min, filtered (0.22 μm filter) and lyophilized. To determine the levels of alginate oxidation and methacrylation, the OMAs were dissolved in deuterium oxide (D_2_O, Sigma) at 2 w/v%, and ^1^H-NMR spectra were recorded on a Varian Unity-300 (300MHz) NMR spectrometer (Varian Inc.) using 3-(trimethylsilyl)propionic acid-d_4_ sodium salt (0.05 w/v%, Sigma) as an internal standard (Figure S20 and Table S1).

### Modification of the Printrbot 3D printer

The 3D printer with a syringe-based extruder was prepared as previously reported. Briefly, the thermoplastic extruder assembly was removed from the 3D printer (Printrbot^®^) and replaced with a custom-built syringe pump extruder (Figure S21). The custom syringe pump extruder was designed to enable the use of the NEMA-17 stepper motor from the original thermoplastic extruder-based printer and mounting of it directly in place of the original extruder on the *x*-axis carriage. The syringe pump extruder was printed with polylactic acid using the thermoplastic extruder on the Printrbot^®^ before its removal. By using the same stepper motor, the syringe pump extruder was natively supported by Cura software (Ultimaker) which is an open source 3D printer host software. The design for the syringe pump extruder and the image files were downloaded as stereolithography (STL) files from the National Institutes of Health 3D Print Exchange (http://3dprint.nih.gov) under an open-source license.

### Ca-crosslinked OMA bioinks

OMAs (2 w/v %) were dissolved in DMEM (Sigma) with a photoinitiator [2-Hydroxy-4’-(2-hydroxyethoxy)-2-methylpropiophenone, 0.05 w/v %, Sigma] at pH 7.4. OMA (1 ml) solutions were loaded into a 1-ml syringe. 40 μl of calcium sulfate slurries (CaSO_4_•2H_2_O) at concentrations of 0, 52.5, 78.8, 91.9, 105.0, and 210.0 mg/ml were added into another 1-ml syringe. After the two syringes were connected together with a female-female luer lock coupler (Value Plastics), the two solutions were mixed back and forth 40 times. The Ca-crosslinked OMA solution was further mixed back and forth 10 times every 10 min for 30 min. Ca-crosslinked OMA bioink was loaded into a 2.5-ml syringe (Gastight Syringe, Hamilton Company) with a 0.5-inch (length) 22 G stainless steel needle (McMaster-Carr). The syringe was then mounted onto the syringe pump extruder on the modified Printrbot 3D printer, the pneumatic extrusion system on a commercial extrusion-based 3D bioprinter (Biobot Basic, Advanced Solutions Life Sciences) or the syringe pump printhead on a commercial syringe pump-based 3D bioprinter (Bio X™, Cellink). The OMA bioinks were printed using Cura software for the modified Printrbot 3D printer, TSIM software for the Biobot Basic printer, or the in-house software in the Bio X™ printer.

For the cell-laden OMA bioinks, human bone marrow-derived mesenchymal stem cells (hMSCs)^2^ were isolated from bone marrow aspirates obtained from the posterior iliac crest of a healthy twenty eight-year old male donor under a protocol approved by the University Hospitals of Cleveland Institutional Review Board. The aspirates were washed with growth medium comprised of DMEM-LG (Sigma) with 10 % prescreened fetal bovine serum (FBS, Gibco). Mononuclear cells were isolated by centrifugation in a Percoll (Sigma) density gradient and the isolated cells were plated at 1.8×10 cells/cm in DMEM-LG containing 10 % FBS and 1 % penicillin/streptomycin (P/S, Thermo Fisher Scientific) in an incubator at 37 °C and 5 % CO_2_. After 4 days of incubation, non-adherent cells were removed and adherent cell were maintained in DMEM-LG containing 10 % FBS, 1 % P/S and 10 ng/ml fibroblast growth factor-2 (FGF-2, R&D) with media changes every 3 days. After 14 days of culture, the cells were passaged at a density of 5×10^3^ cells/cm^2^, cultured for an additional 14 days, and then stored in cryopreservation media in liquid nitrogen until use. To encapsulate hMSCs into OMA bioink, hMSCs were expanded in growth media consisting of DMEM-LG with 10% FBS (Sigma), 1 % P/S and 10 ng/ml FGF-2. To prepare the cell-laden OMA bioinks, hMSCs (passage 3) were harvested with trypsin/EDTA (Thermo Fisher) and concentrated by centrifugation at 300 × g for 5 min. Following aspiration of the supernatant, pelleted hMSCs were suspened in OMA solution (5×10^6^ cell/ml), and then the OMA solution with suspended cells was crosslinked as described above.

### Rheological properties of OMA bioinks

Dynamic rheological examination of the OMA bioinks was performed to evaluate their mechanical properties and shear-thinning, shear-yielding and self-healing behavior with a Kinexus ultra+ rheometer (Malvern Panalytical). In oscillatory mode, a parallel plate (25 mm diameter) geometry measuring system was employed, and the gap was set to 1 mm. After each OMA bioink was placed between the plates, all the tests were started at 25 ± 0.1 °C, and the plate temperature was maintained at 25 °C. To determine the shear-thinning and shear-yielding behaviors of the OMA bioinks, viscosity change was measured as a function of shear rate and shear stress, respectively. Oscillatory frequency sweep (0.1-10 Hz at 1 % shear strain) tests were performed to measure storage moduli (G’) and loss modulis (G”). Oscillatory strain sweep (0.10-100 % shear strain at 1 Hz) tests were performed to determine the G’/G” crossover. To demonstrate the self-healing properties of the OMA bioinks, cyclic deformation tests were performed at 100 % shear strain with recovery at 1 % shear strain, each for 1 min at 1 Hz.

### 3D printing of OMA bioinks

OMA or hMSC-laden OMA bioinks were loaded into syringes as described above, connected to 0.5-inch (length) 22 G stainless steel needles and mounted on the Printrbot 3D printer. The tip of each needle was positioned at the center and 0.1 mm from the bottom of the glass dish (Cellink), and the print instructions were sent to the printer using Cura software. For 3D printing of patterned structures, bioink-loaded syringes were connected to a 0.5-inch (length) 22 G stainless needles and mounted onto the commercial extrusion-based 3D bioprinter (Biobot Basic) or the commercial syringe pump-based 3D bioprinter (Bio X’, Cellink), and the bioink was printed using the software indicated earlier for these printers.

### Analysis of 3D printed structures

Linear filaments of the OMA bioink were printed with 22, 25 and 27 G needles using the BioX™ printer, photocrosslinked under UV light a 20 mW/cm for 1 min. and then filaments were imaged using a fluorescence microscope (TMS-F, Nikon) equipped with a digital camera (Coolpix 995, Nikon). Diameters of the 3D OMA filaments were measured at least 400 times using 10 different printed filaments for each group using ImageJ (National Institutes of Health).

To assess the quality and accuracy of the 3D printing of OMA bioinks, cuboids [10×10×5 mm, Figure 3(c)] were printed with 20 and 22 G needles. The dimensions were then measured using a caliper. The measured dimensions were compared to the original dimensions specified in the 3D models. The fidelity (%) was calculated as Length_measured_/Length_original_ × 100 (N=3).

### Cytotoxicity of 3D printing process

Viability of hMSCs in the photocrosslinked 3D printed constructs was investigated using Live/Dead staining comprised of fluorescein diacetate (FDA, Sigma) and ethidium bromide (EB, Fisher Scientific). The staining solution was freshly prepared by mixing 1 ml FDA solution (1.5 mg/ml in dimethyl sulfoxide, Sigma) and 0.5 ml EB solution (1 mg/ml in PBS, Thermo Fisher Scientific) with 0.3 ml PBS (pH 8). After photocrosslinking of 3D printed constructs (5 mm diameter and 1 mm height) in a 24-well tissue culture plate under UV light at 20 mW/cm^2^ for 1 min, 1 ml growth media was added into each well. 20 μl of staining solution was added into each well and incubated for 3-5 min at room temperature, and then stained hMSC-hydrogel constructs were imaged using a fluorescence microscope (ECLIPSE TE 300) equipped with a digital camera (MU1402-NI05, AmScope).

### Chondrogenesis

After 3D printing of the bioinks, the 3D printed constructs were further stabilized by photocrosslinking under UV at 20 mW/cm^2^ for 1 min. After photocrosslinking, 3D printed constructs were transferred into 100 ml spinner flasks (Bellco Glass Inc., Vineland, NJ) containing 80 ml of chondrogenic differentiation media [1 % ITS+ Premix, 100 nM dexamethasone, 37.5 μg/ml l-ascorbic acid-2-phosphate, 1 mM sodium pyruvate, 100 μM nonessential amino acids, and 10 ng/ml TGF-β_1_ in HG-DMEM] or growth media as a control. The spinner flasks were placed in a humidified incubator at 37 °C with 5% CO_2_ and stirred at 40 rpm. The chondrogenic media was changed every other day. After 4 weeks of culture, 3D printed hMSC constructs were harvested, fixed in 10 % neutral buffered formalin overnight at 4 °C. Whole intact 3D printed ears were stained with Toluidine blue O. Fixed chondrogenically differentiated and control 3D printed cuboids (3×3×1 mm) were embedded in paraffin, sectioned at a thickness of 10 μm, stained with Toluidine blue O, and then imaged using a microscope (TMS-F, Nikon) equipped with a digital camera (Coolpix 995, Nikon). To measure GAG production, chondrogenically differentiated 3D printed cuboids (3×3×1 mm) were homogenized at 35000 rpm for 60 sec using a TH homogenizer (Omni International) in buffer (1 mL, pH 6.5) containing papain (25 μg m/l, Sigma), l-cysteine (2 × 10^-3^M, Sigma), sodium phosphate (50 × 10^-3^M, Thermo Fisher Scientific) and EDTA (2×10^-3^M, Thermo Fisher Scientific) and then digested at 65 °C overnight. GAG content was quantified by a dimethylmethylene blue assay and DNA content was measured using the PicoGreen^®^ assay. GAG content was normalized to DNA content.

### Statistical analysis

Quantitative data were expressed as mean ± standard deviation. Statistical analysis was performed by one-way analysis of variance (ANOVA) with the Tukey significant difference post hoc test using Prism software (GraphPad). A value of p<0.05 was considered statically significant.

## Acknowledgements

The authors gratefully acknowledge funding from the National Institutes of Health’s National Institute of Arthritis and Musculoskeletal and Skin Diseases under award numbers R01AR069564 and R01AR066193. The contents of this publication are solely the responsibility of the authors and do not necessarily represent the official views of the National Institutes of Health.

## Data availability

The datasets generated and/or analyzed during the current study are available from the corresponding author upon request.

## Author contributions

**O. J.:** Conceptualization, Methodology, Formal analysis, Investigation, Writing-Original draft, Review & Editing. **Y.B.L.:** Formal analysis, Investigation, Writing-Original draft, Review & Editing. **S.J.L.:** Investigation, Writing-Original draft, Review & Editing. **N.G.:** Investigation, Writing-Original draft, Review & Editing. **J.L.:** Investigation, Writing-Original draft, Review & Editing. **E.A.:** Supervision, Conceptualization, Writing-Original draft, Review & Editing.

