## Supplemental figures and table for "Stem cell-laden hydrogel bioink for generation of high resolution and fidelity engineered tissues with complex geometries"

**Supporting Figures**

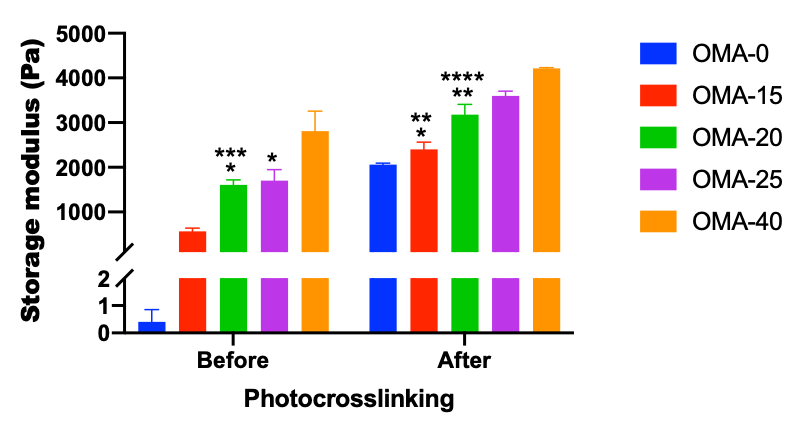

**Figure S1**. Storage moduli of the OMA (1OX20MA) bioinks synthesized from high viscosity alginate before and after photocrosslinking. *p>0.05 compared to OMA-0 after photocrosslinking. **p>0.05 compared to OMA-40 before photocrosslinking. ***p>0.05 compared to OMA-25 before photocrosslinking. ****p>0.05 compared to OMA-25 after photocrosslinking. Otherwise, p<0.05.

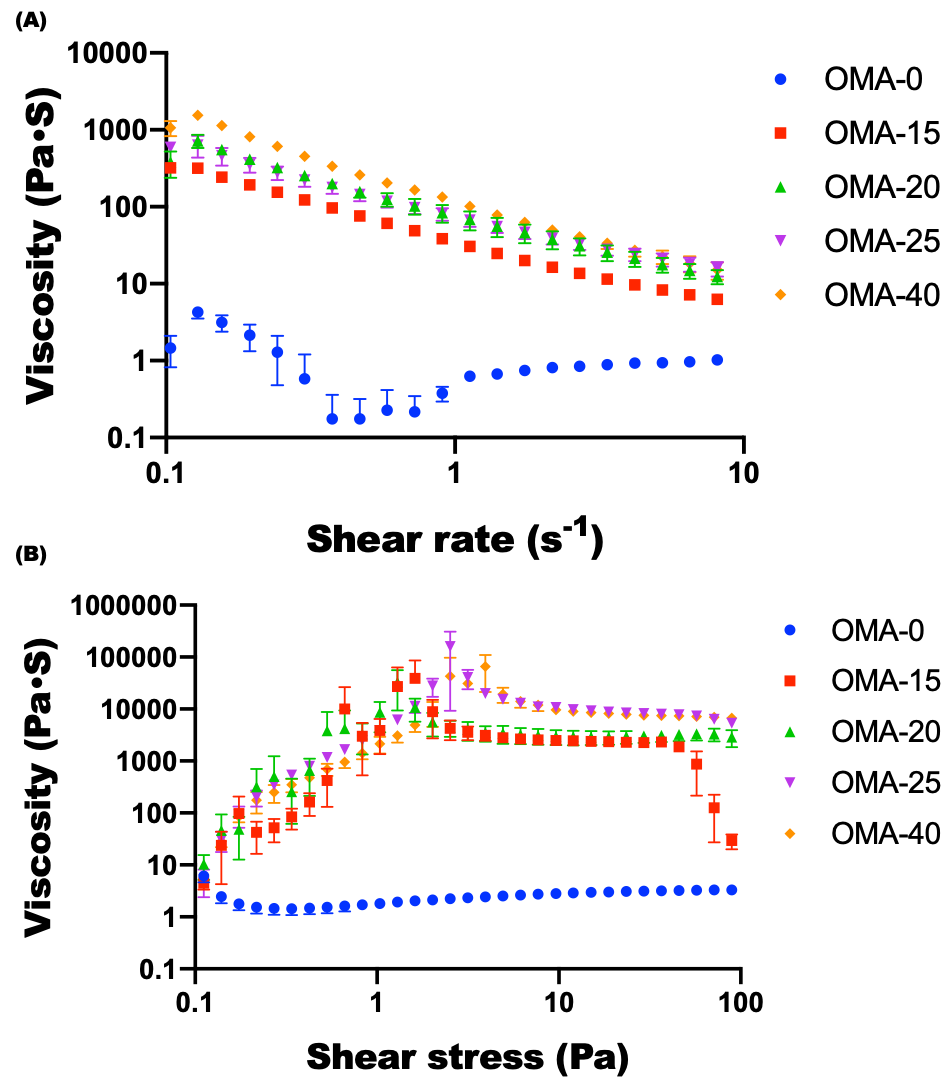

**Figure S2.** Viscosity of the OMA (2OX20MA; 2 and 20 % theoretical oxidation and methacrylation, respectively) bioinks synthesized from high viscosity alginate as a function of (A) shear rate and (B) shear stress.

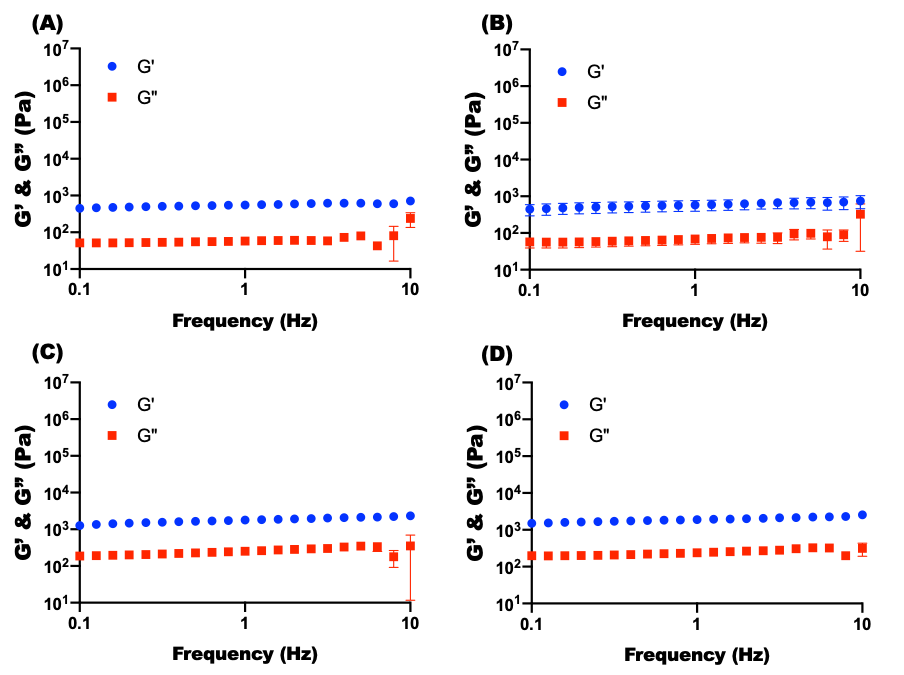

**Figure S3.** Frequency sweep tests of the (A) OMA-15, (B) OMA-20, (C) OMA-25 and (D) OMA-40 bioinks prepared with the 2OX20MA OMA synthesized from high viscosity alginate.

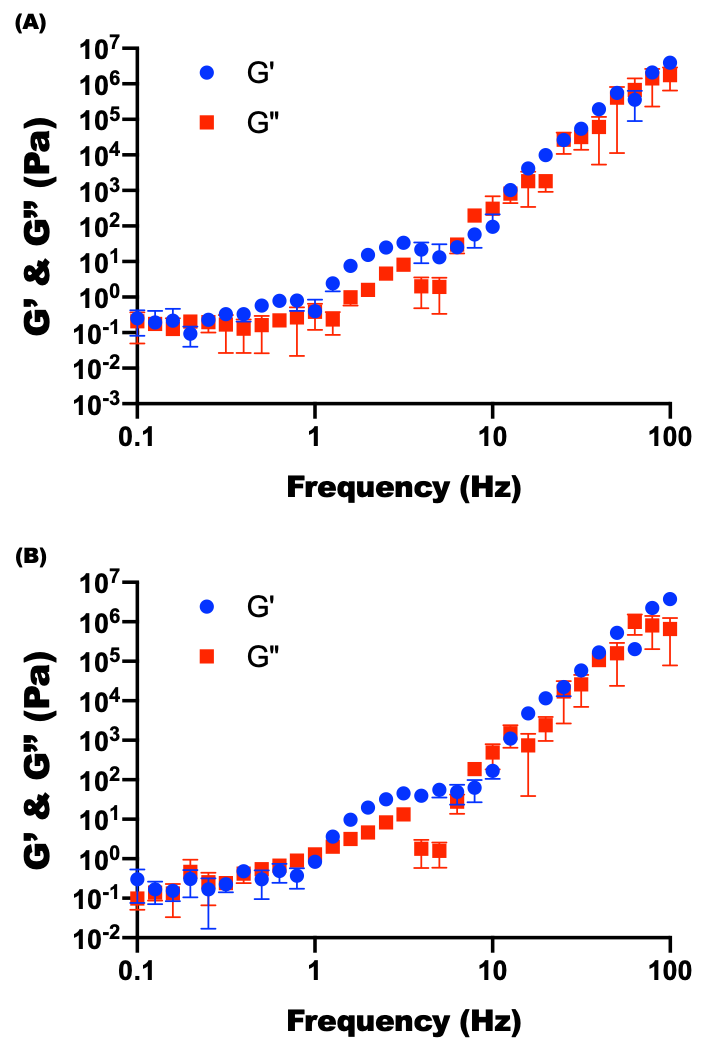

**Figure S4.** Frequency sweep tests of OMA-0 with the (A) 1OX20MA and (B) 2OX20MA OMAs synthesized from high viscosity alginate.

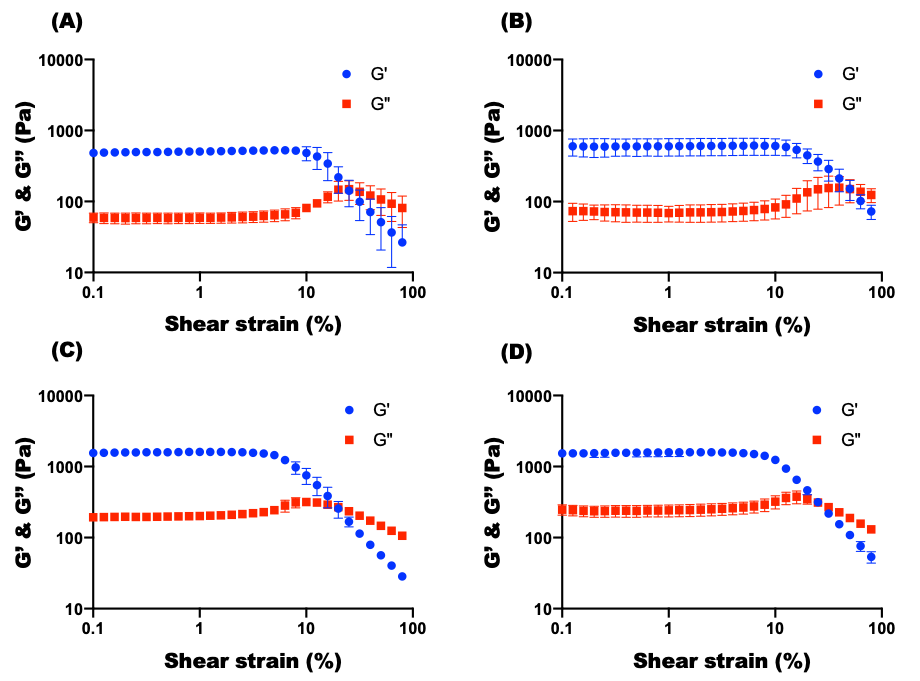

**Figure S5.** Strain sweep tests of the (A) OMA-15, (B) OMA-20, (C) OMA-25 and (D) OMA-40 bioinks prepared with the 2OX20MA OMA synthesized from high viscosity alginate.

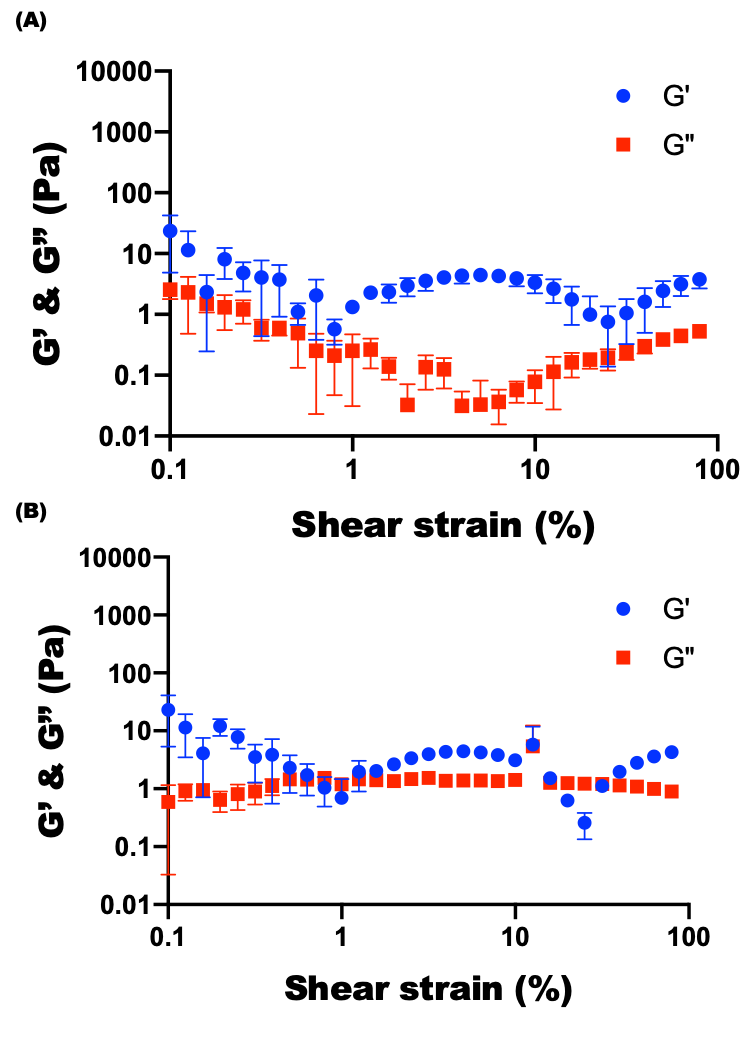

**Figure S6.** Strain sweep tests of the OMA-0 prepared with (A) 1OX20MA and (B) 2OX2OMA OMAs synthesized from high viscosity alginate.

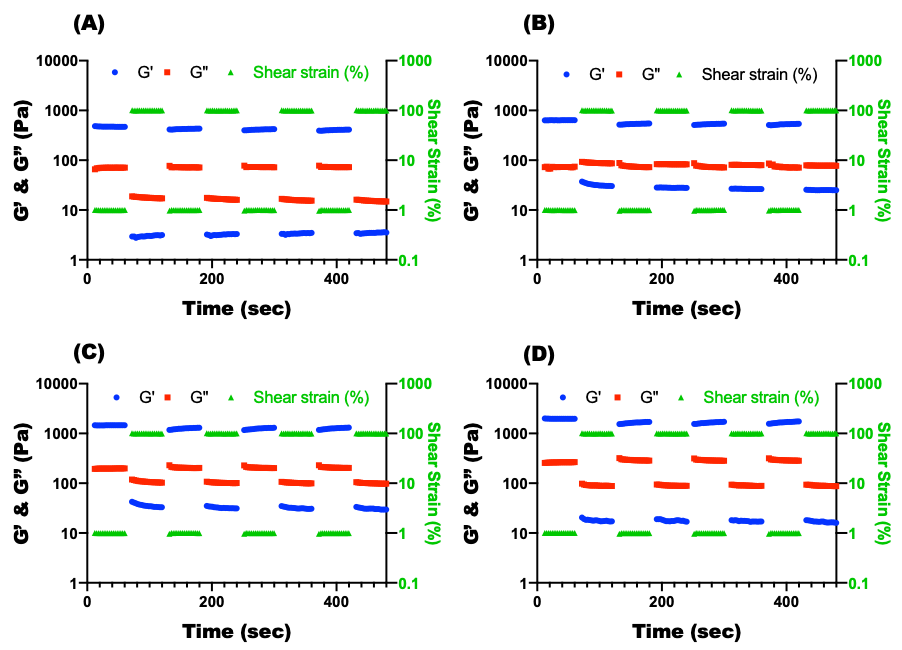

**Figure S7.** Shear moduli changes in dynamic strain tests of the (A) OMA-15, (B) OMA-20, (C) OMA-25 and (D) OMA-40 bioinks prepared with the 2OX20MA synthesized from high viscosity alginate with alternating low (1 %) and high (100 %) strains at 1 Hz.

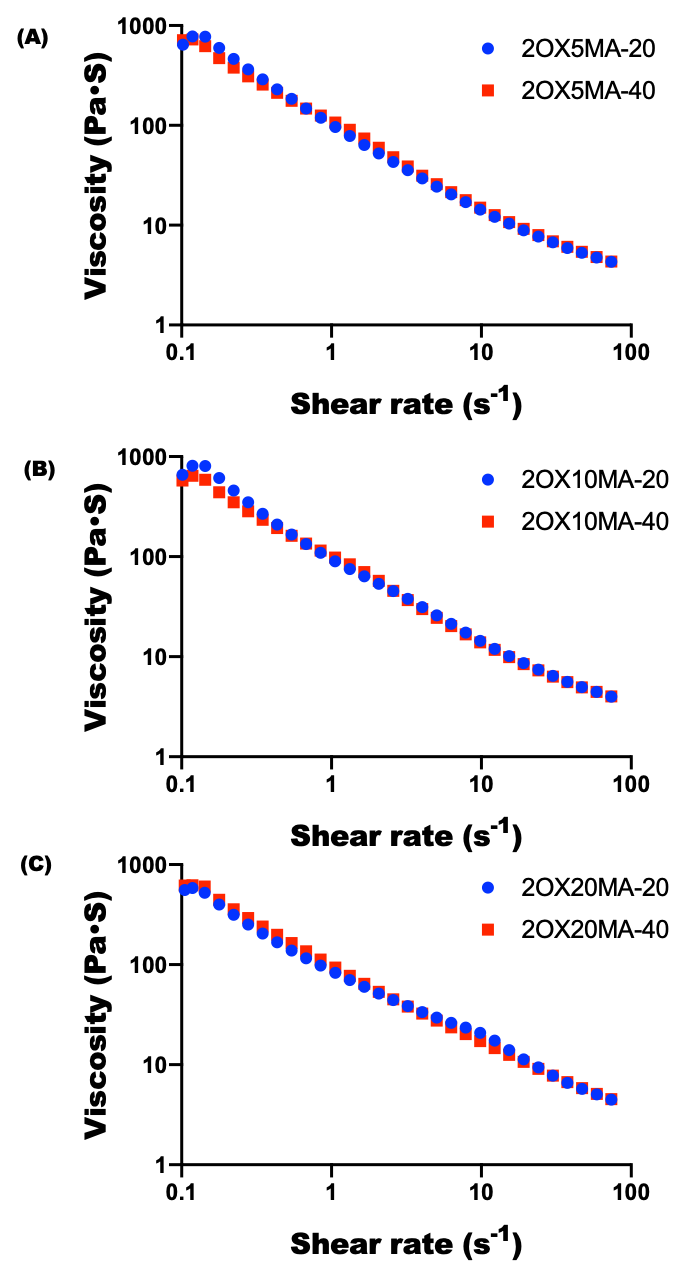

**Figure S8.** Viscosity as a function of shear rate of the OMA bioinks prepared with (A) 2OX5MA (2 and 5 % theoretical oxidation and methacrylation, respectively), (B) 2OX10MA (2 and 10 % theoretical oxidation and methacrylation, respectively), and (C) 2OX20MA (2 and 20 % theoretical oxidation and methacrylation, respectively) synthesized from low viscosity alginate and mixed with 20 or 40 μl CaSO_4_ (1.22 M).

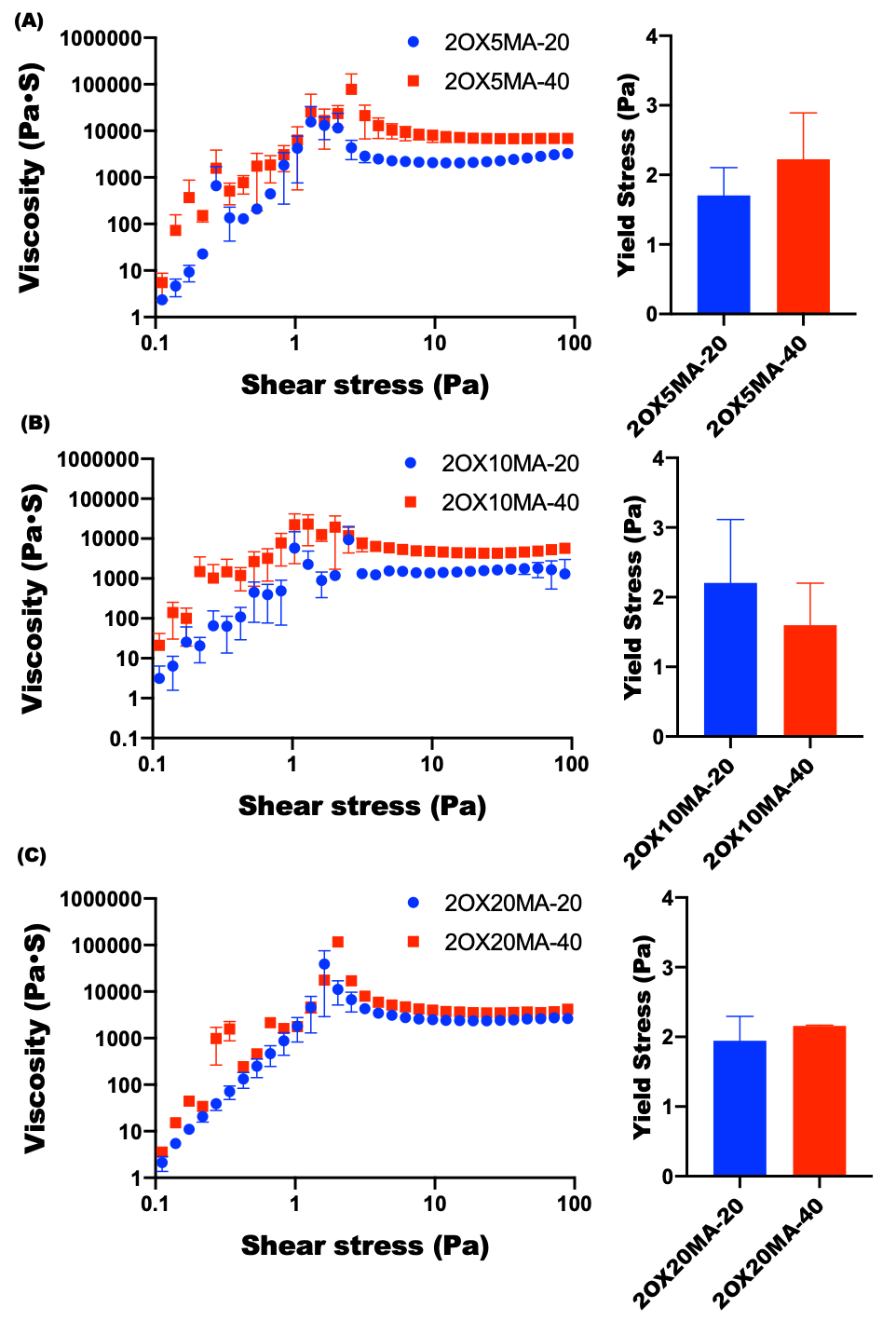

**Figure S9.** Viscosity measurements as a function of shear stress and yield stress of bioinks composed of (A) 2OX5MA, (B) 2OX10MA and (C) 2OX20MA low viscosity OMAs and 20 or 40 μl CaSO_4_ (1.22 M).

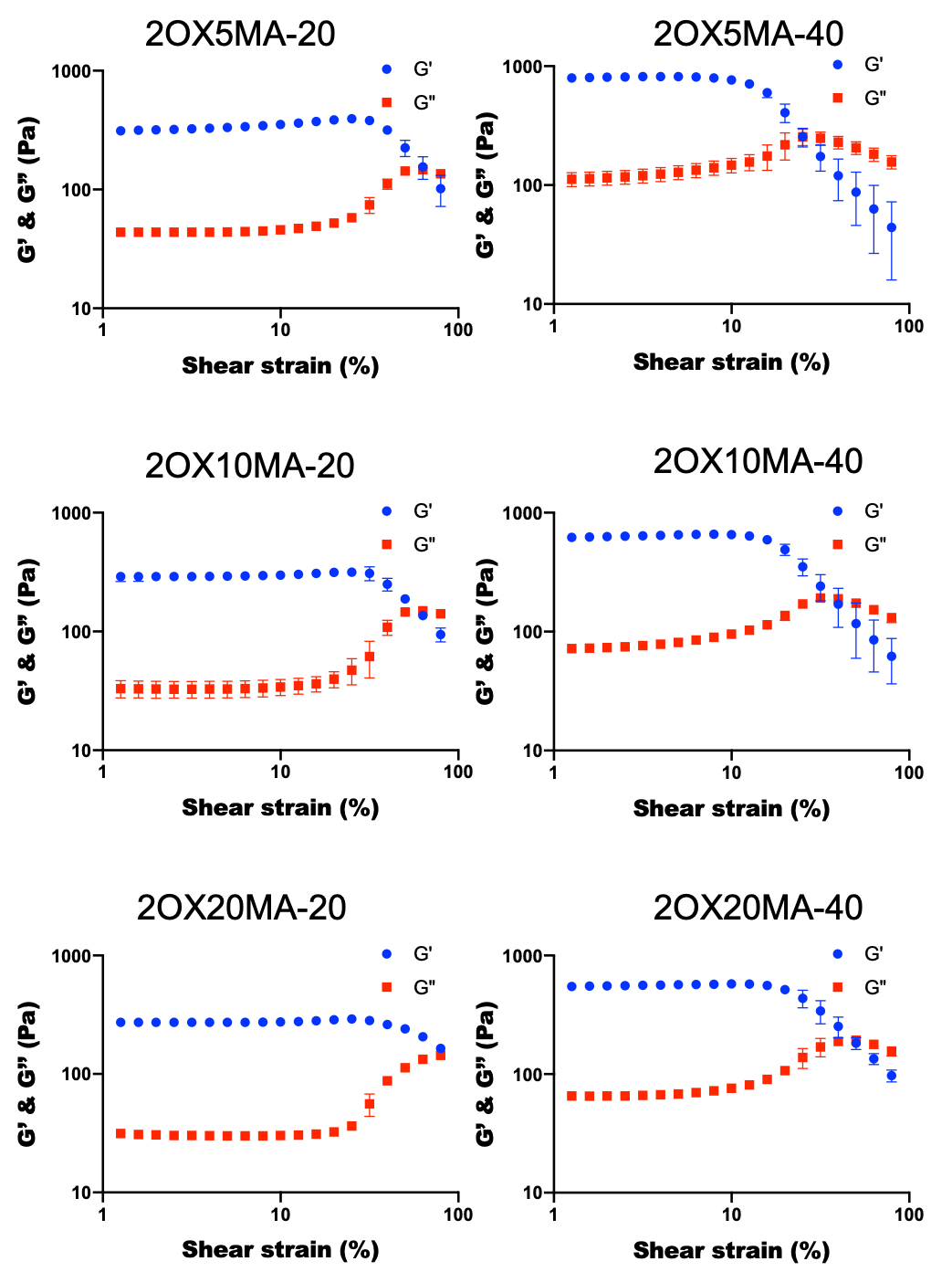

**Figure S10.** Strain sweep tests of OMA bioinks synthesized from low viscosity alginate.

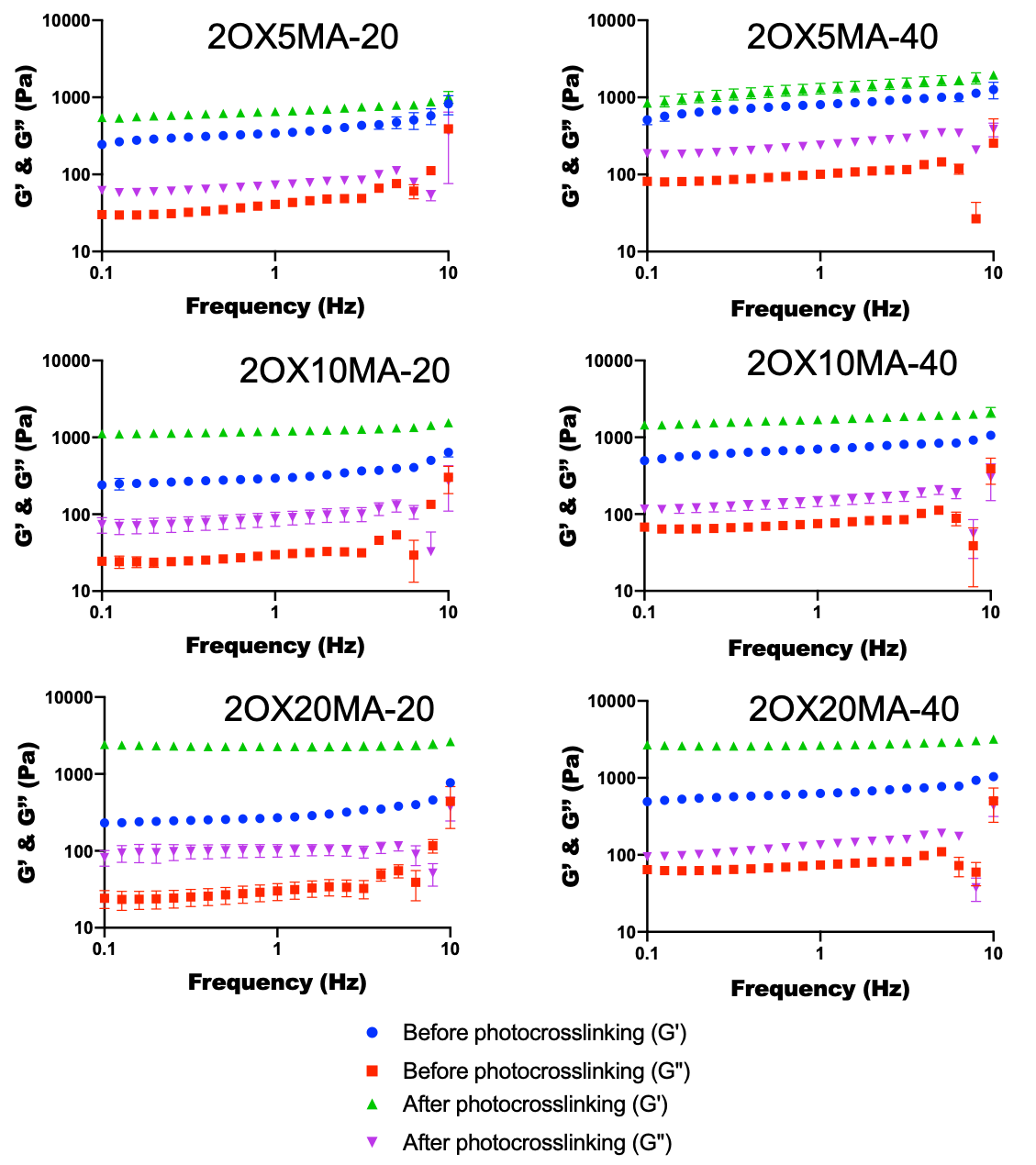

**Figure S11.** Frequency sweep tests of OMA bioinks synthesized from low viscosity alginate before and after photocrosslinking.

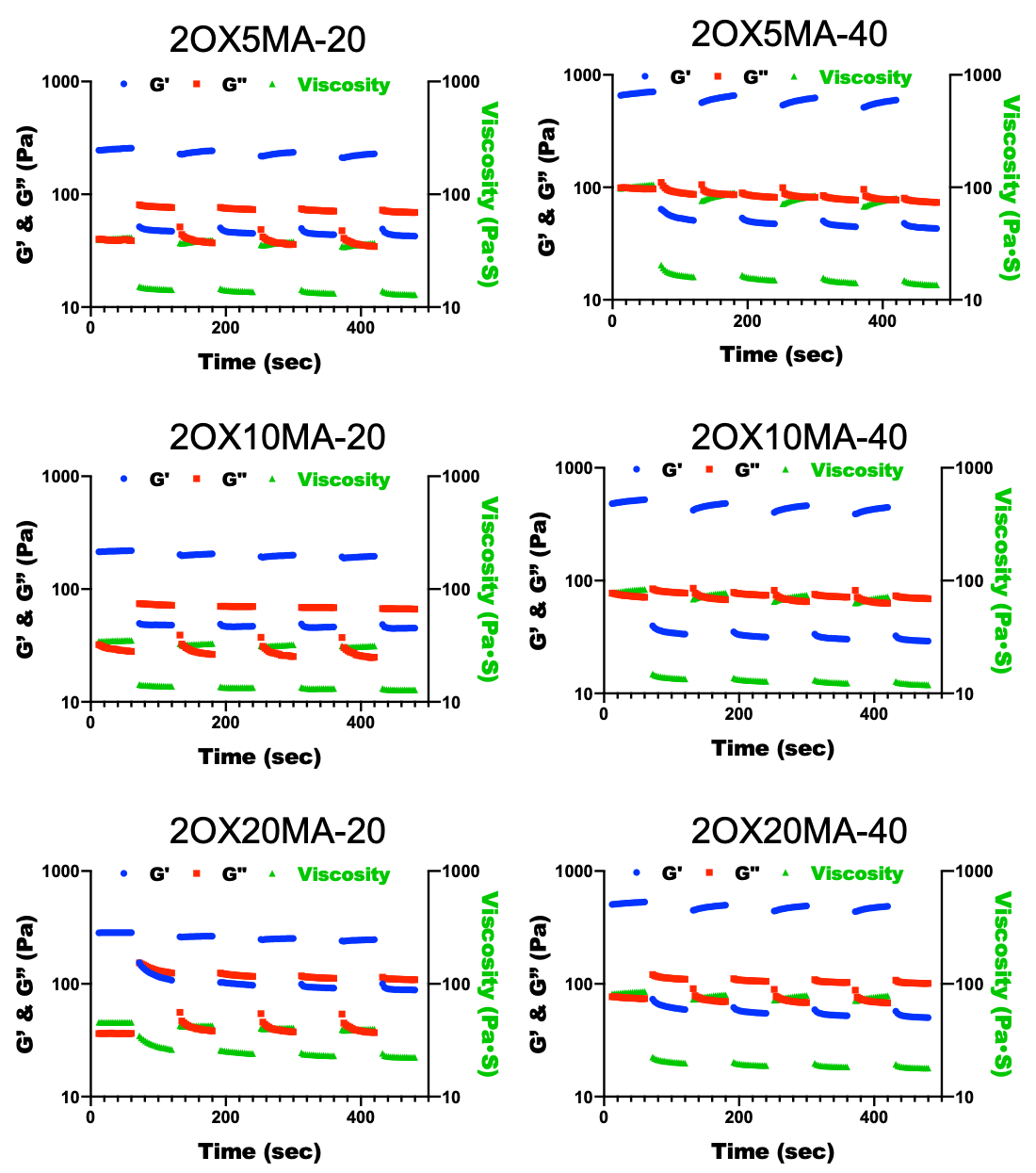

**Figure S12.** Shear moduli and viscosity changes in dynamic strain tests of the OMA bioinks synthesized from low viscosity alginate with alternating low (1 %) and high (100 %) strains at 1 Hz.

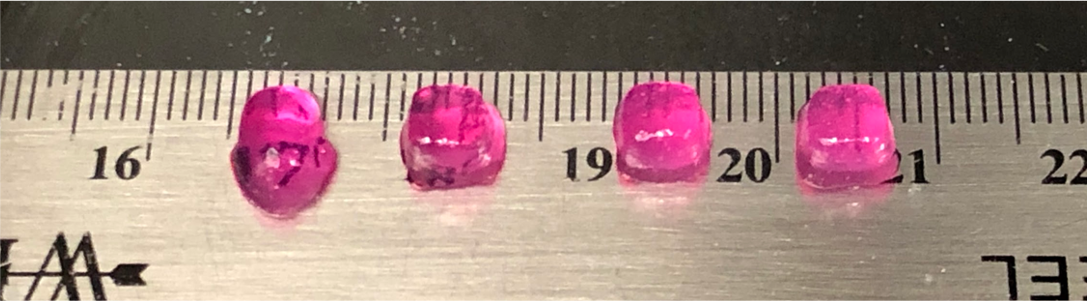

**Figure S13.** Photographs of the 3D printed structures using the BioX printer and OMA bioinks synthesized from high viscosity 2OX20MA OMA and.

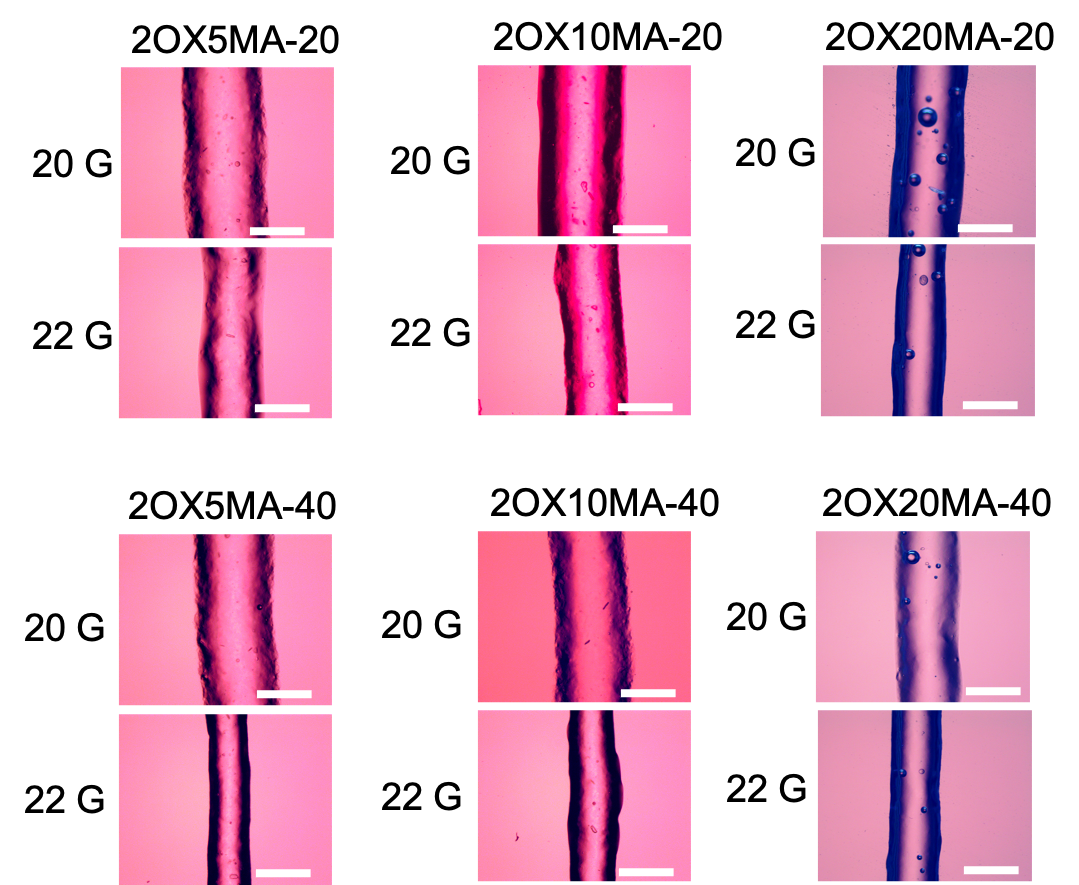

**Figure S14.** Photomicrographs of OMA bioinks synthesized from low viscosity alginate 3D printed in the form of straight line filaments with 20 and 22 G printing needles. The scale bars indicate 500 μm.

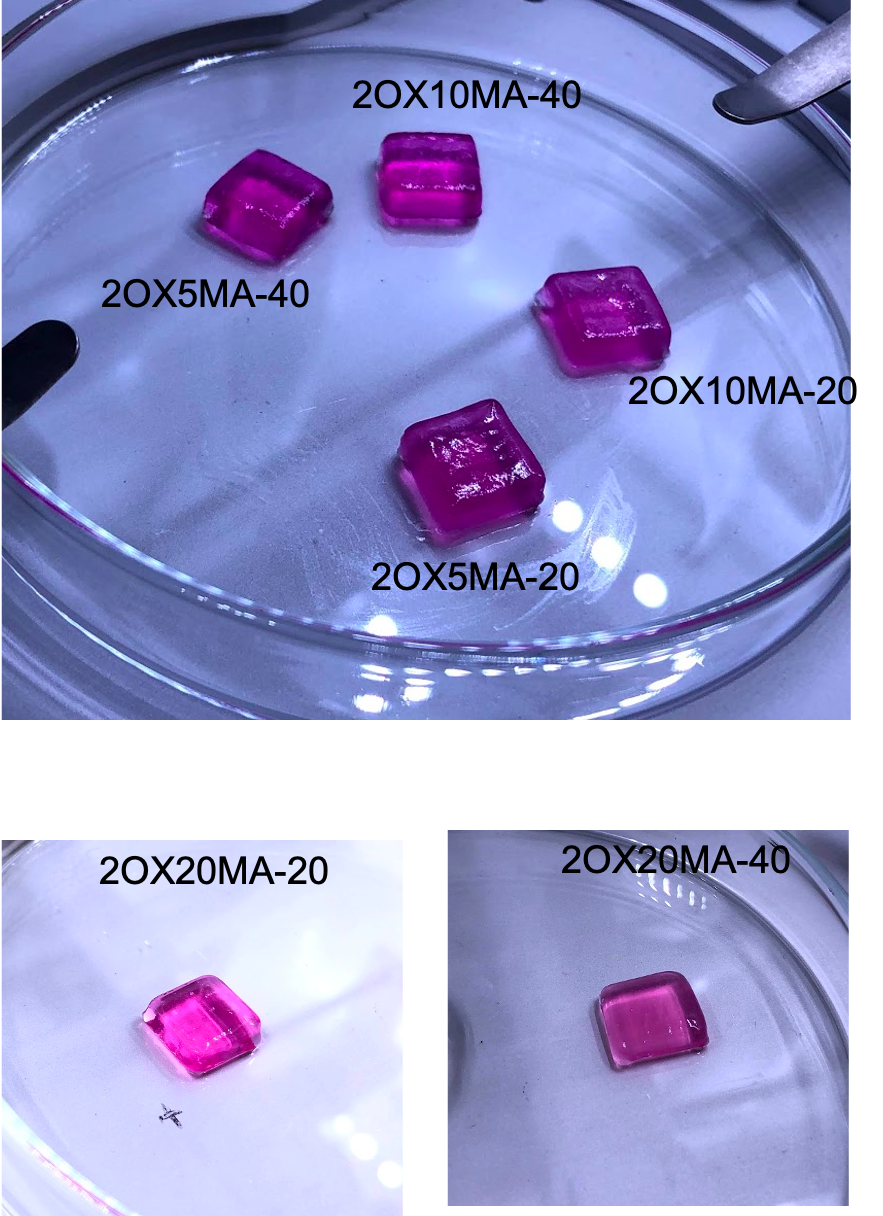

**Figure S15.** Photographs of the 3D printed structures (10×10×5 mm) using the BioX printer and OMA bioinks synthesized from low viscosity alginate.

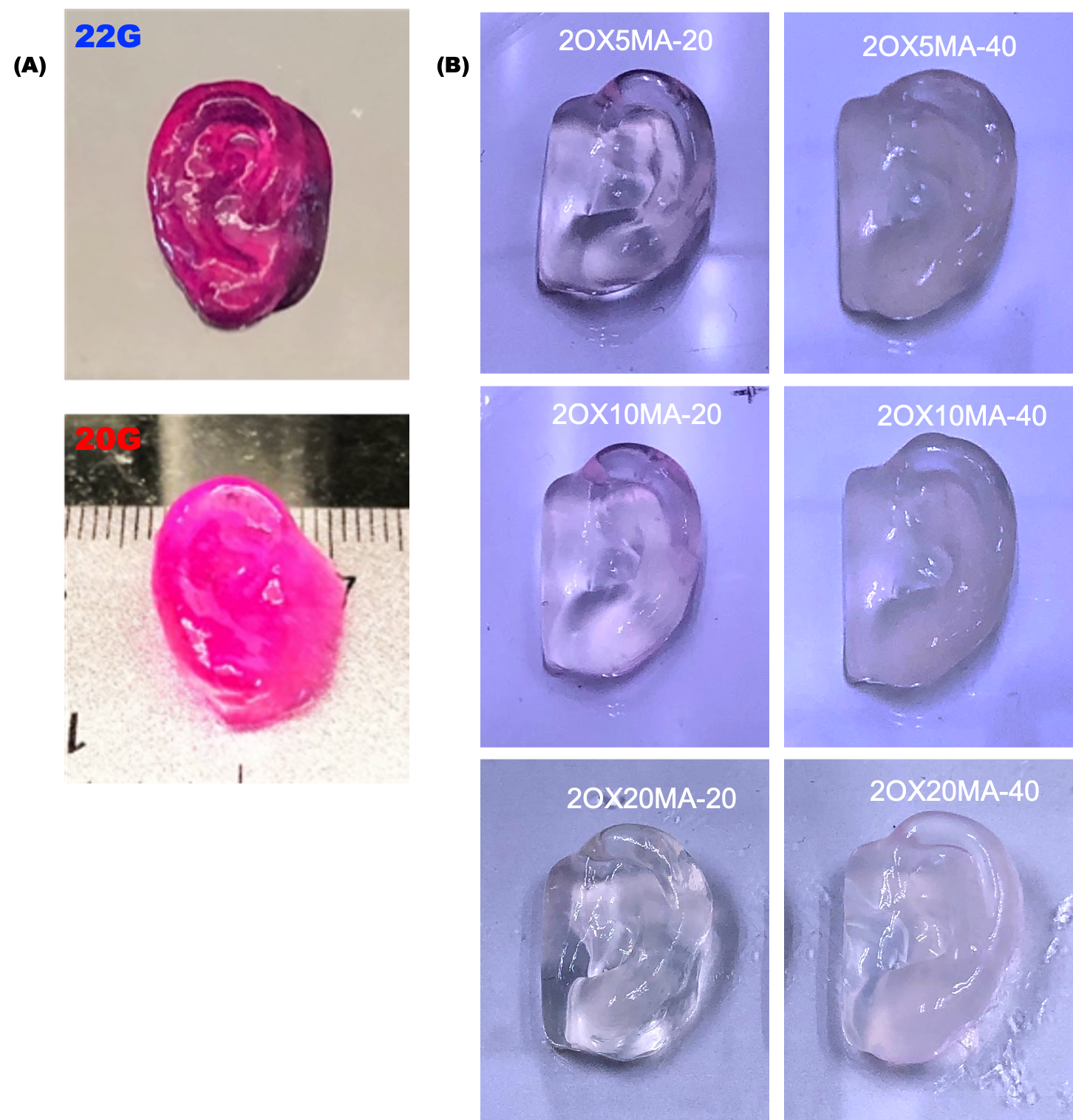

**Figure S16.** Photographs of ears 3D printed on the BioX printer with (A) 22 and 20 G printing needles using 1OX20MA synthesized from high viscosity alginate and mixed with 20 μl CaSO_4_ (1.22 M) and (B) a 22 G printing needed using different OMA formulations synthesized from low viscosity alginate.

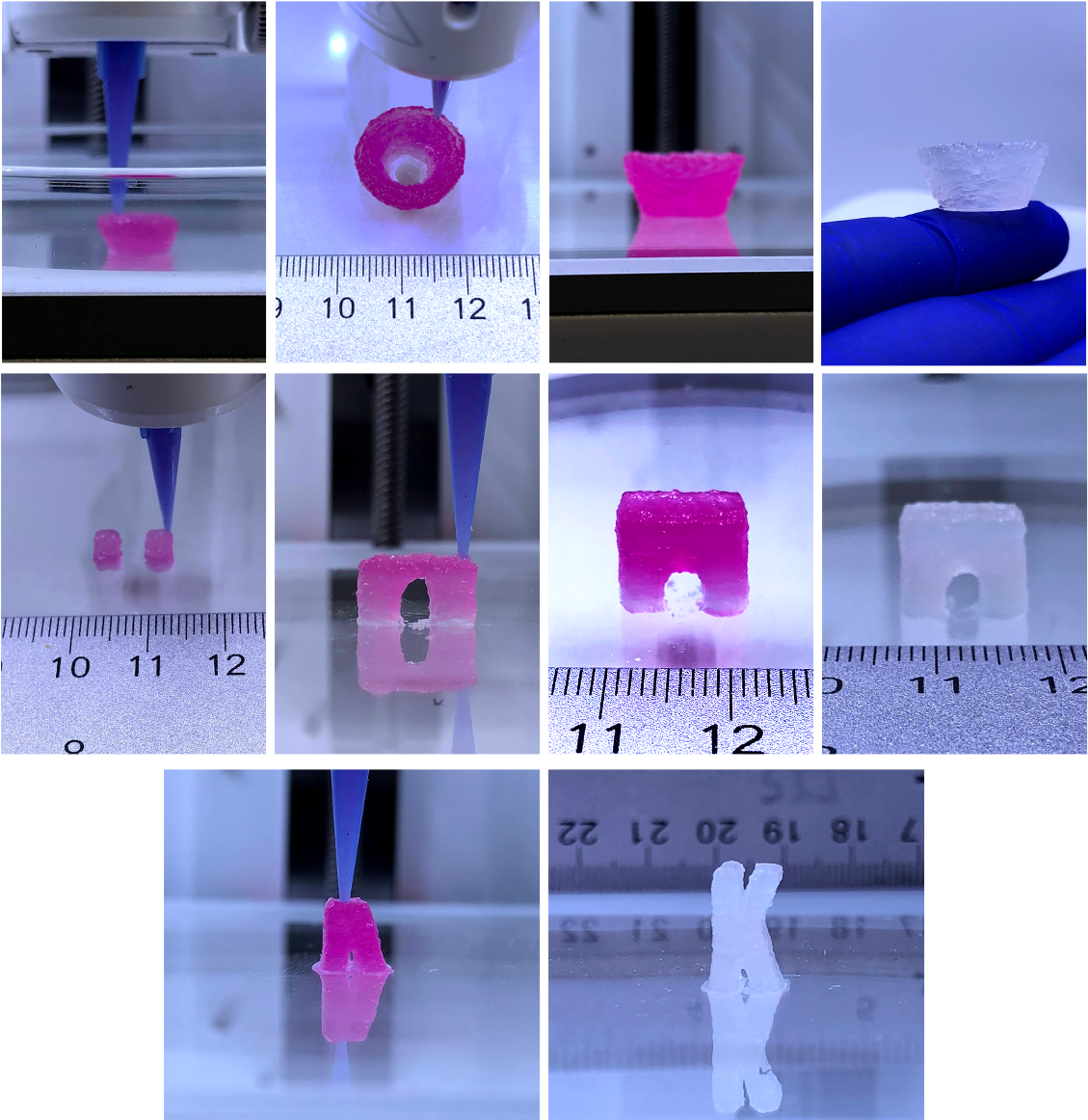

**Figure S17.** Photographic images, which were taken during printing and after printing and photocrosslinking, of the overhang geometries 3D (i.e., a bowl, bridge, the letter “K”) printed on the BioX printer with a 22 G printing nozzle using the OMA bioink (1OX20MA) synthesized from high viscosity alginate and mixed with 40 μl CaSO_4_ (1.22 M).

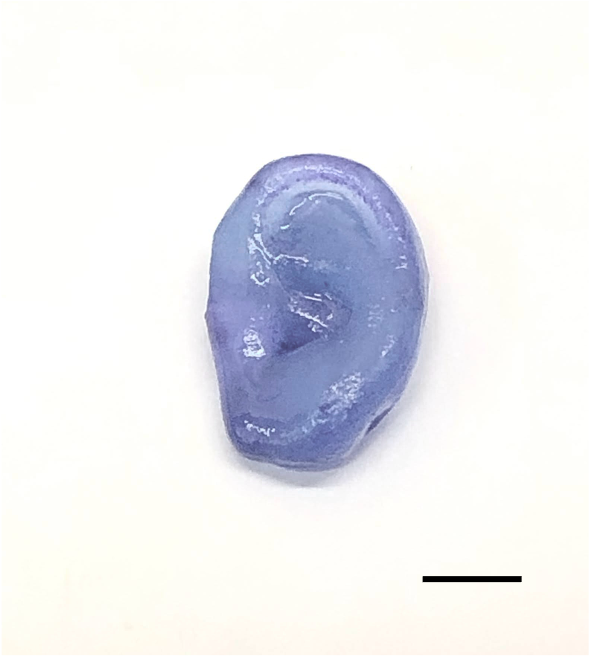

**Figure S18.** Toluidine blue O stained photographic image of a 3D printed ear, which was printed using the modified Printrbot printer with hMSC-laden OMA bioink synthesized from high viscosity alginate and cultured in growth media for 4 weeks. The scale bar indicates 1 cm.

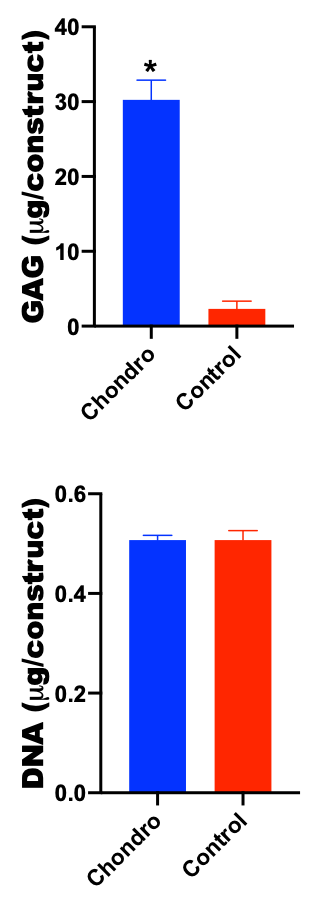

**Figure S19.** Quantification of the GAG and DNA contents in the chondrogenically differentiated 3D printed constructs. *p<0.05 compared to Control.

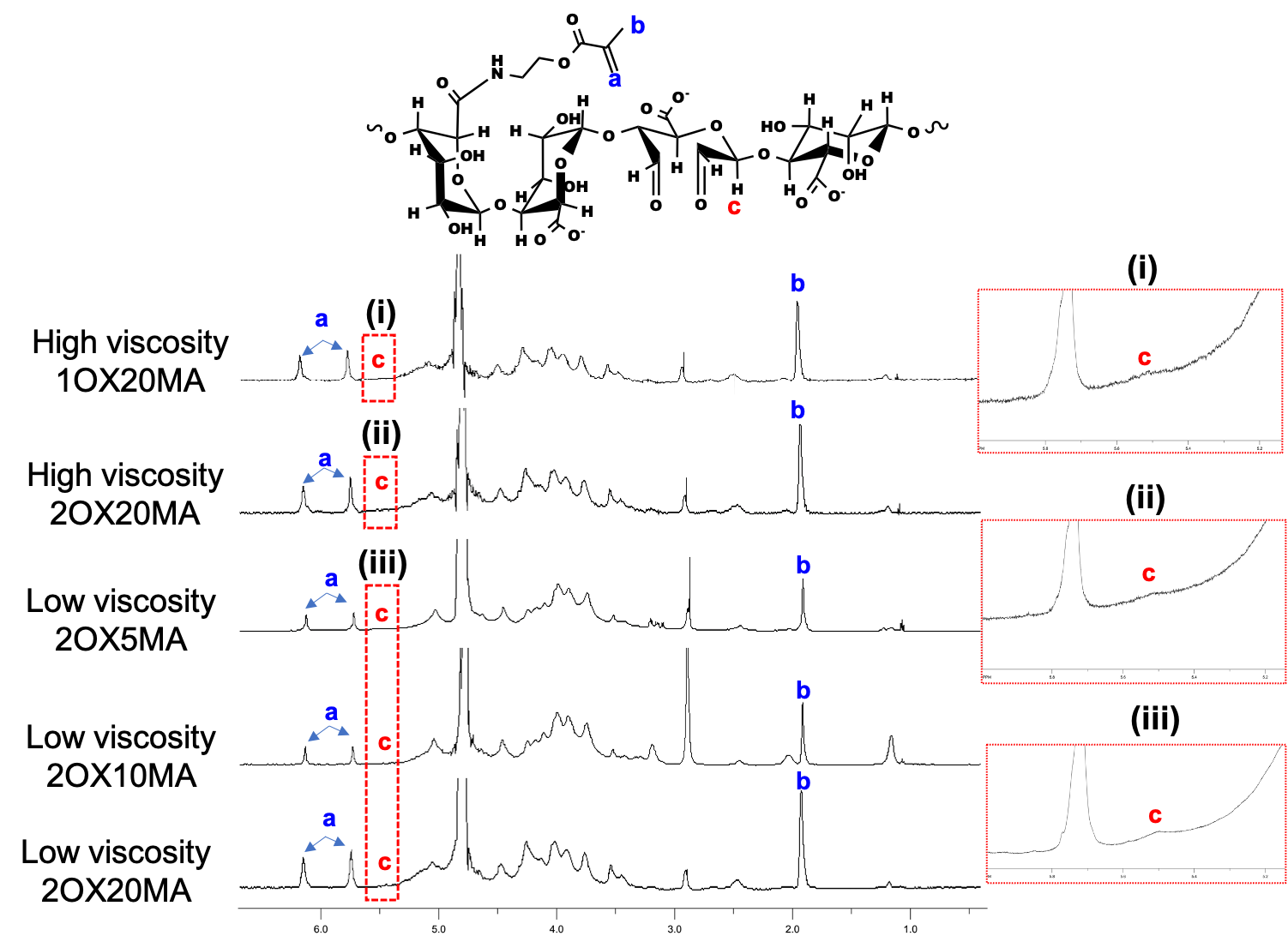

**Figure S20.** ^1^H-NMR spectra of OMAs with various degrees of oxidation and methacrylation in D_2_O.

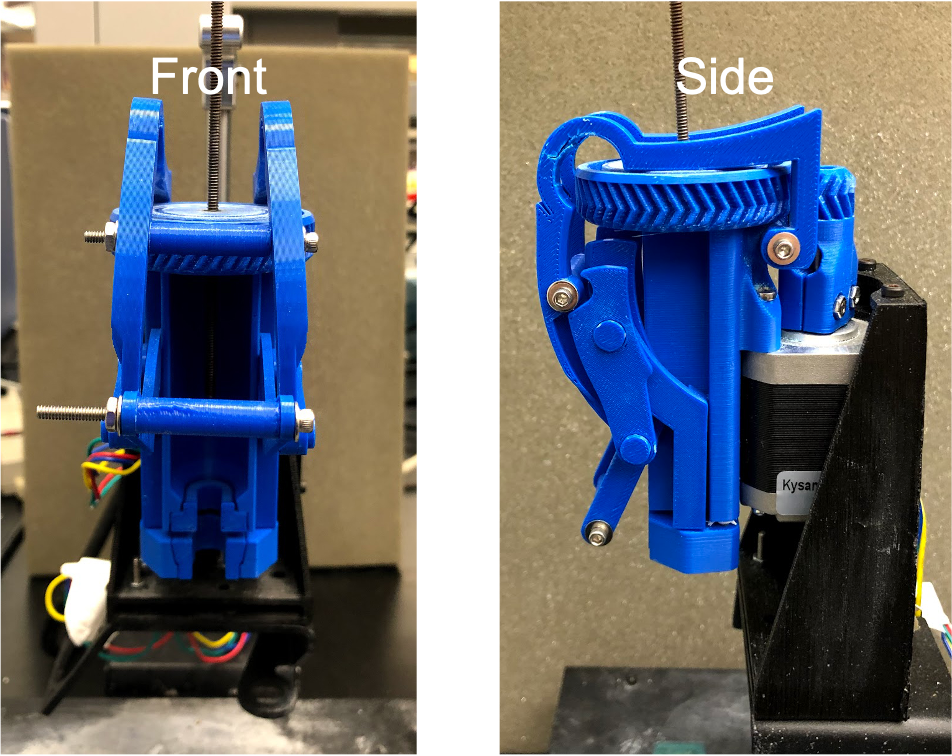

**Figure S21.** The syringe pump extruder mounted on the modified Printrbot 3D printer.

**Table S1.** Actual oxidation (%) and methacrylation (%) of OMAs.

| Code | Alginate | Theoretical oxidation (%) | Theoretical methacrylation (%) | Actual oxidation (%) | Actual methacrylation (%) |
| --- | --- | --- | --- | --- | --- |
| 1OX20MA | High viscosity LF120M | 1 | 20 | 0.85 | 10.61 |
| 2OX20MA | High viscosity LF120M | 2 | 20 | 1.78 | 11.94 |
| 2OX5MA | Low viscosity LF120M | 2 | 5 | 1.91 | 4.68 |
| 2OX10MA | Low viscosity LF120M | 2 | 10 |  | 8.43 |
| 2OX20MA | Low viscosity LF120M | 2 | 20 |  | 13.39 |
